# Root-knot nematodes produce functional mimics of tyrosine-sulfated plant peptides

**DOI:** 10.1101/2022.10.13.511487

**Authors:** Henok Zemene Yimer, Dee Dee Luu, Alison Coomer Blundell, Maria Florencia Ercoli, Paulo Vieira, Valerie M. Williamson, Pamela C. Ronald, Shahid Siddique

## Abstract

Root-knot nematodes (*Meloidogyne* spp.) are highly evolved obligate parasites that threaten global food security. These parasites have a remarkable ability to establish elaborate feeding sites in roots, which are their only source of nutrients throughout their life cycle. A wide range of nematode effectors have been implicated in modulation of host pathways for feeding site development. Plants produce a diverse array of peptide hormones including plant peptides containing sulfated tyrosine (PSYs), which promote root growth via cell expansion and proliferation. A sulfated PSY-like peptide RaxX (required for activation of XA21 mediated immunity X) produced by the biotrophic bacterial pathogen (*Xanthomonas oryzae* pv. *oryzae*), has been previously shown to contribute to bacterial virulence. Here, we report the identification of genes from root-knot nematodes predicted to encode PSY-like peptides (MigPSYs) with high sequence similarity to both bacterial RaxX and plant PSYs. Sulfated synthetic peptides corresponding to predicted MigPSYs stimulate root growth in Arabidopsis. *MigPSY* transcript levels are highest early in the infection cycle. Down-regulation of *MigPSY* gene expression reduces root galling and egg production, suggesting that the MigPSYs serve as nematode virulence factors. Together these results indicate that nematodes and bacteria utilize similar sulfated peptides to hijack plant developmental signaling pathways to facilitate parasitism.

## Introduction

Plant-parasitic nematodes (PPNs) are among the most destructive plant pathogens, causing an annual economic loss of $8 billion to U.S. growers and over $100 billion worldwide (Nicol, 2011, Sasser & Freckman, 1986). Root-infecting sedentary endoparasitic nematodes including root-knot nematodes (RKNs; *Meloidogyne* spp.) and cyst nematodes (*Heterodera* spp. and *Globodera* spp.) cause the greatest economic damage. The RKN species *M. incognita, M. javanica*, and *M. arenaria* boast extremely broad host ranges and can infect thousands of species, including annual and perennial crops and both dicots and monocots (Trudgill & Blok, 2001). Infective second-stage juveniles (J2) of RKNs invade the plant close to the root tip. Once inside the root, J2s migrate between cells until they reach the vascular tissue (Holbein et al., 2019), where they become immobile and induce the formation of several highly modified, adjacent multinucleated giant cells. While giant cells develop, neighboring cells hypertrophy and divide, leading to the formation of characteristic galls (root-knots) (Rutter et al., 2022, Williamson & Gleason, 2003). Cell walls of giant cells undergo intensive modification that includes both thickening and loosening to allow expansion and support nutrient uptake. These feeding sites are the only source of nutrients for nematodes throughout their approximately one-month long life cycle (Siddique et al., 2022).

The establishment of a long-term feeding site allows PPNs to acquire nutrients from their plant host and is essential for their development and reproduction. A wide range of nematode-secreted molecules have been implicated in establishing and maintaining these feeding sites. Among these molecules are small peptides that resemble plant peptide hormones. The best studied are mimics of CLE (CLAVATA/embryo surrounding region) peptides in cyst nematodes. In that group of PPNs, CLE-like peptides are delivered as propeptides into the host cytoplasm through the nematode stylet. The propeptides are then post-translationally modified and proteolytically processed by host enzymes for secretion into the host’s extracellular space as mature CLE-like peptides. The N-terminal variable domain of the propeptide is required for this process (Guo et al., 2011, Wang et al., 2010, Wang et al., 2021). Nematode CLEs are thought to facilitate the formation of host feeding sites by activating CLE signaling pathways (Gheysen & Mitchum, 2019, Mitchum & Liu, 2022). RKNs also encode mimics of other peptide hormones including CEP (C-terminally encoded peptides), RALF (rapid alkalization factors), and IDA (inflorescence deficient in abscission) peptides (Gheysen & Mitchum, 2019, Zhang et al., 2020, Mitchum & Liu, 2022, Bobay et al., 2013, Kim et al., 2018, Tucker & Yang, 2013, Wang et al., 2005). In contrast to cyst nematode CLEs, all of the RKN peptide hormone mimics characterized so far lack a variable domain and may therefore be directly delivered as mature peptides to the host’s extracellular space (Gheysen & Mitchum, 2019).

Plant peptides containing sulfated tyrosine (PSYs) promote root growth via cell expansion and proliferation (Amano et al., 2007, Tost et al., 2021). PSY1 was identified from Arabidopsis suspension cell culture medium as an 18-amino acid tyrosine sulfated glycopeptide derived from a 75-amino acid precursor peptide (Amano et al., 2007). PSY peptides have been identified in all higher plants and more recently in mosses and bacteria (Tost et al., 2021). Tyrosine sulfation is catalyzed by tyrosylprotein sulfotransferase (TPST), which is encoded by a single gene in Arabidopsis (At1g08030) (Komori et al., 2009). Exogenous application of AtPSY1 to Arabidopsis seedlings activates plasma membrane localized proton pumps and promotes acidification of the extracellular space (Fuglsang et al., 2014). This acidification leads to pH-dependent activation of expansins and other cell wall modifying enzymes that loosen and soften the cell wall (Cosgrove, 2000).

Several *Xanthomonas* species secrete a tyrosine sulfated peptide RaxX (required for activation of XA21-mediated immunity X) with sequence similarity to a 13-amino acid motif conserved throughout plant PSYs (Pruitt et al., 2017). RaxX peptides from diverse species of *Xanthomonas* mimic the growth-promoting activities of PSYs (Liu et al., 2019). The proposed role of PSY peptides in cell expansion and proliferation led us to hypothesize that PPNs might also produce and secrete PSY-like peptides into host plants to facilitate formation of the nematode feeding site. Here, we identify RKN genes encoding several PSY-like peptides with high sequence similarity to both bacterial RaxX and plant PSYs. We further characterize their expression and investigate their potential roles in parasitism.

## Results and Discussion

### Root-knot nematode genes encode PSY-like peptides

We used the conserved 13-amino acid sequence PSY motif of RaxX (RaxX13^-C^) to interrogate predicted translation products of cDNA sequences from available databases (Blanc-Mathieu et al., 2017, Bolt et al., 2018). This conserved motif contains a tyrosine residue immediately preceded by an aspartic acid and followed by a NXXHXP sequence of 6 residues downstream (**Figure 1A**). The aspartic acid-tyrosine residue pair is the minimum requirement for tyrosine sulfation (Stone et al., 2009), a post-translational modification critical for PSY activity (Amano et al., 2007). Following re-interrogation with initial hits, we identified 11 cDNA translation products (**Table S1**) in the closely related RKN species *Meloidogyne incognita, M. javanica*, and *M. arenaria*. As *M. incognita, M. javanica*, and *M. arenaria* are closely related asexual species that belong to the *Meloidogyne incognita* group, we will refer to the nematode genes as *MigPSY*s (*Meloidogyne incognita* group PSYs) thereafter.

**Figure 1.**
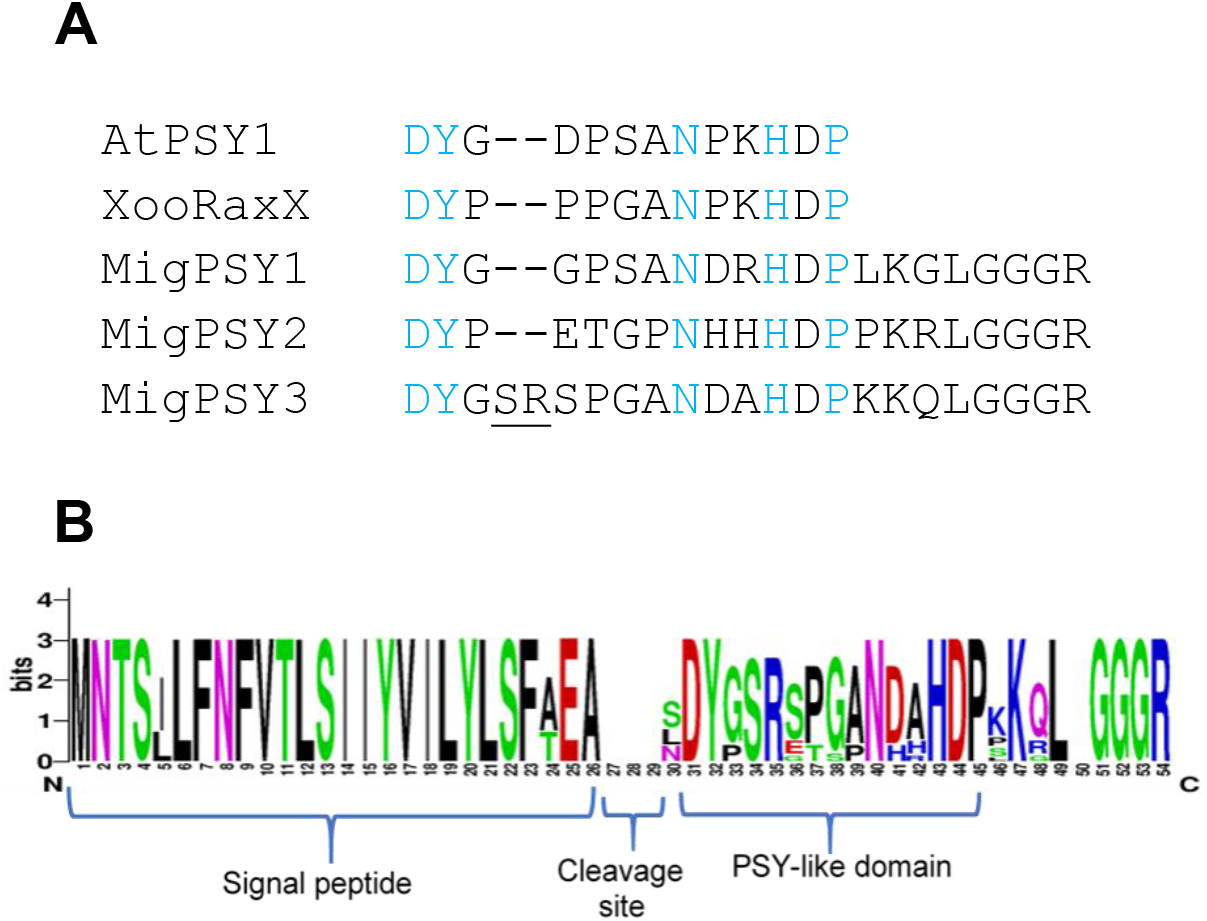
Root-knot nematodes are predicted to encode PSY-like peptides. **(A)** Amino acid sequences of the conserved PSY motif from Arabidopsis [*Arabidopsis thaliana*] and from the bacterial species, Xoo (RaxX), compared with those from the three MigPSYs. Letters in blue represent the conserved amino acids of the PSY motif. For each of the MigPSY types the predicted product extending form the conserved motif to the predicted C-terminus is shown. **(B)** Protein logo alignment of 11 predicted *MigPSY* gene products predicted in the species *M. incognita, M. javanica*, and *M. arenaria* (Table S2). The primary translation products are predicted to contain a highly conserved N-terminal secretion signal peptide followed by a PSY-like domain beginning 1-4 amino acids after the signal peptide cleavage site.

Based on differences in amino acid residues within the conserved domain, we classified MigPSYs into three types: MigPSY1, MigPSY2, and MigPSY3 (**Figure 1A**). We identified MigPSY1 in only one isolate of *M. arenaria*, while MigPSY2 was present in *M. arenaria* and *M. javanica* but not in *M. incognita*. For MigPSY3, two amino acids (SR) were inserted between the conserved DY and NXXHXP motifs. MigPSY3s were the most widely represented, with 2–3 copies present in the genome of each MIG species. We also identified closely related PSY-like motifs in genome drafts of additional Clade I *Meloidogyne* species (*M. enterolobii, M. floridensis*, and *M. luci*), but not in more distantly related *Meloidogyn*e spp. or in other groups of PPNs.

All identified *MigPSYs* are predicted to produce primary translation products of 50– 51 amino acids (**Figure 1B**). Each is predicted to be cleaved after a highly conserved N-terminal signal peptide, resulting in a 21–24 amino acid peptide with the conserved PSY motif at or very close to its N-terminus. No transmembrane domain was predicted by DeepTMHMM analysis (Jeppe Hallgren et al., 2022). Without additional processing, each translation product would contain a C-terminal GGGR sequence (**Table S2, Fig. 1B**). For all 11 genes, two exons are spliced together upon transcript processing to constitute the aspartate residue that initiates the PSY-like domain. MigPSYs, like other RKN peptide mimics (IDA, CEP, RALF, CLE), lack a variable domain and may therefore be directly delivered to the host’s extracellular space.

### Exogenous application of MigPSY peptides enhances plant root growth

Based on our bioinformatic analysis, we hypothesized that nematode peptides would be secreted and that these peptides would have PSY peptide-like activity. To test this hypothesis, we measured the growth response of Arabidopsis roots after peptide treatment. For these experiments, we utilized a loss-of-function mutant in Arabidopsis tyrosyl protein sulfotransferase (*tpst-1*), which is required for tyrosine sulfation, a crucial post-translational modification for PSY activity. The *tpst-1* mutant displays a stunted root phenotype (Komori et al., 2009) that can be partially rescued by exogenous treatment with sulfated PSY or RaxX peptides (Pruitt et al., 2017). We assessed the root growth response of *tpst-1* mutant seedlings to treatment with synthetic peptides corresponding to the predicted mature peptide sequence of each of the three MigPSY types or with the previously characterized peptides RaxX21 (a 21-amino acid derivative of RaxX from *Xanthomonas oryzae* pv. *oryzae*) and AtPSY1 as controls (Pruitt et al., 2017). All peptides were sulfated at their N-terminal tyrosine (**Table S3**). We grew seedlings on Murashige and Skoog (MS) medium containing individual synthetic MigPSY peptides provided at a concentration of 100 nM and measured root lengths nine days after sowing. Each of the MigPSY peptides significantly increased the root length of *tpst-1* Arabidopsis seedlings to an extent similar to plants treated with the control peptides AtPSY1 and RaxX21 (**Figure 2**). RaxX13, which contains only the conserved 13-amino acid PSY-like domain (denoted by ^-C^), is sufficient for RaxX to induce root growth (Pruitt et al., 2017). Similarly, truncated MigPSY peptides containing only the conserved DYX_5-7_NXXHXP PSY-like domain (MigPSY^-C^) also promoted *tpst-1* root growth. However, root lengths were only moderately increased with the truncated MigPSY3^-C^ peptide treatment compared to treatment with the complete mature MigPSY3 peptide.

**Figure 2.**
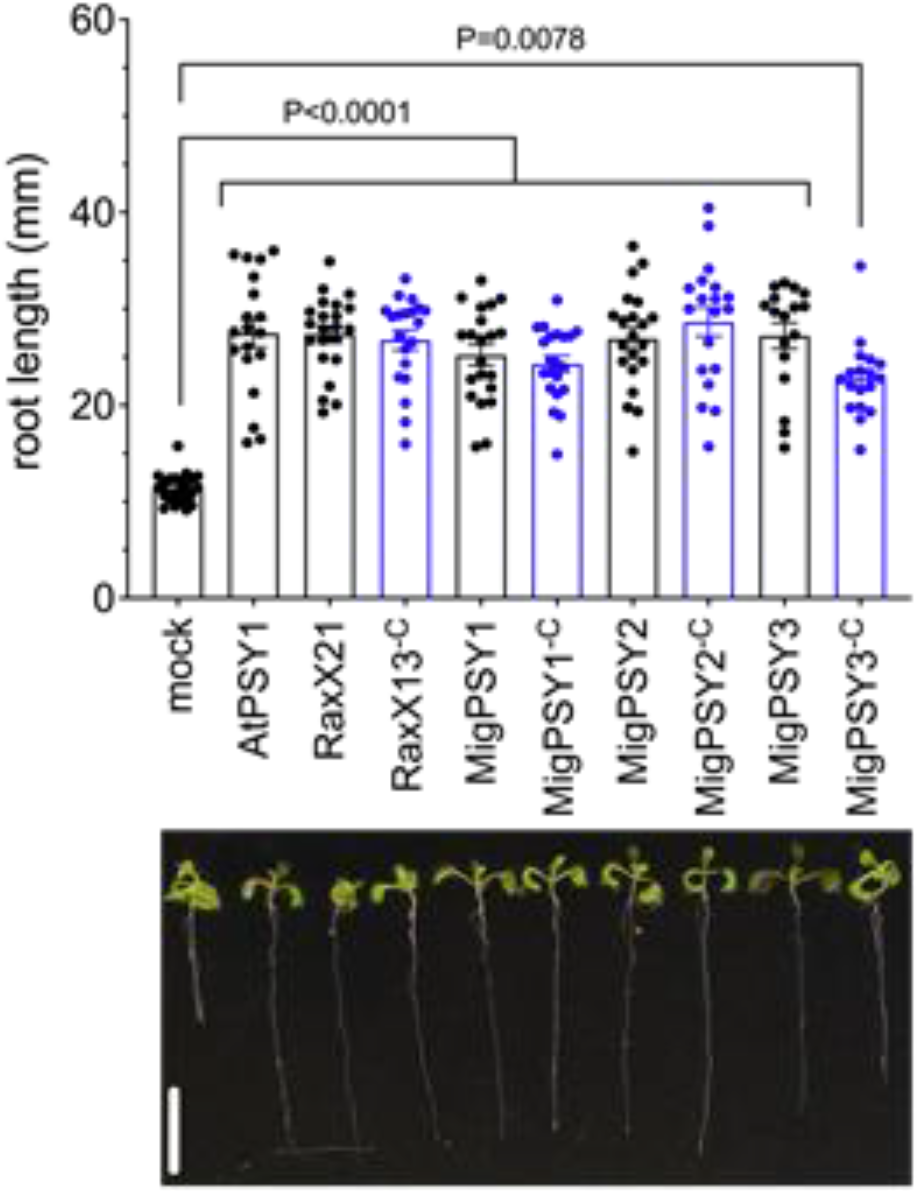
Exogenous treatment with MigPSY peptides promotes root growth. Arabidopsis *tpst-1* mutant seeds were sown on MS media containing either water (mock) or 100 nM of synthetic tyrosine sulfated peptide derived from the predicted MigPSY mature sequence (black) or just the conserved DYX_5-7_NXXHXP PSY-like sequence (denoted by ^-C^, blue). AtPSY1 from Arabidopsis and RaxX21 and RaxX13 from *Xoo* were used as positive controls. Bars represent the mean root length (mm) ± SEM of at least 17 seedlings, shown as individual dots (top panel). Statistical significance of peptide-treated seedlings to mock-treated seedlings were analyzed using the Kruskal-Wallis test. Representative seedlings are shown in the bottom panel (scale bar: 1 cm). Data represent one of three independent experiments with similar results.

### *MigPSY* gene expression is highest at early stages of nematode infection

We analyzed the relative expression levels of *MigPSY* in pre-parasitic juveniles and during infection. For this, we used *M. javanica* (Lambert et al., 1999). *M. javanica* can infect over a thousand host species, including tomato (*Solanum lycopersicum*) and rice (*Oryza sativa*). Database searches indicated that the *M. javanica* genome harbors one copy of *MigPSY2* and two closely related copies of *MigPSY3*, but does not encode *MigPSY1* (**Table S1;** Szitenberg et al., 2017). To investigate the expression pattern of *MigPSYs* during host infection, we designed qPCR primers that can efficiently amplify both *MigPSY2* and *MigPSY3* (**Supplementary Figure 1)**. We collected several hundred root segments containing galls from tomato and rice plants inoculated with *M. javanica* at four time points after inoculation (2, 4, 8, and 21 days after inoculation [dai]). We selected these time points to represent the following stages of parasitism: entry into the root and migration to the feeding site (2 dai); the initial formation of giant cells and galls (4 dai); third molt (J3) nematodes (8 dai); and final molt into adult females (21 dai). Compared to their expression levels in infective juveniles, *MigPSY* expression levels were higher at 2 and 4 dai in both tomato and rice plant roots (**Figure 3**). However, *MigPSY* expression decreased at 8 dai and was barely detectable at 21 dai in both hosts. The induced expression of *MigPSYs* at the early stages of nematode infection suggests that their gene products may be involved in facilitating migration within the root and/or developing nematode feeding sites.

**Figure 3.**
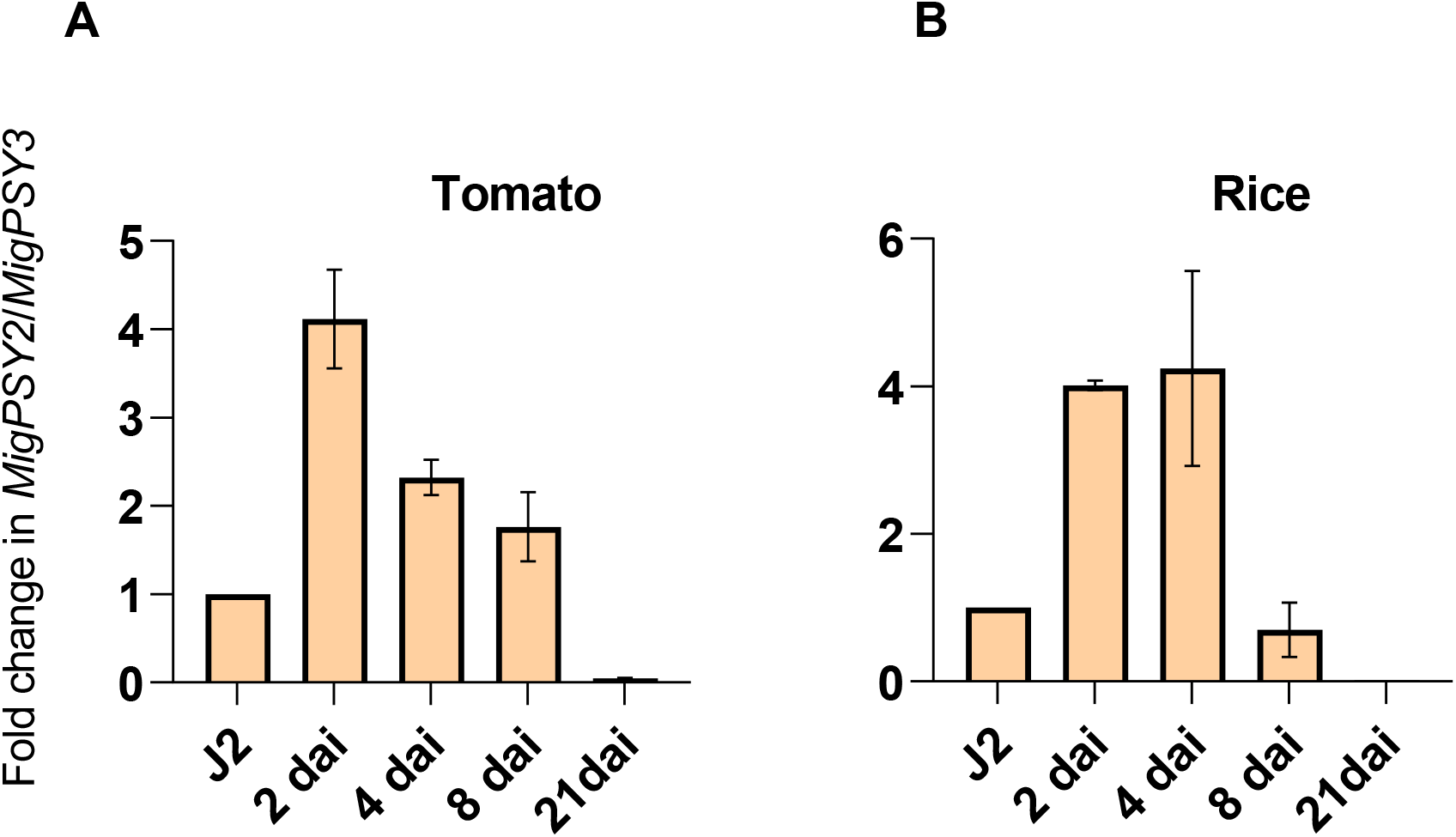
*MigPSY* expression is induced during early stages of nematode infection. Relative expression levels of *MigPSY* genes in *M. javanica* following infection of tomato (A) or rice (B) roots at the indicated time points after infection, as determined by reverse transcription quantitative PCR (RT-qPCR). Values represent relative expression levels with the level in infective second-stage juveniles (J2s) set to 1. The mRNA levels were measured as three technical replicates per sample. In (A), the transcript levels of each gene were normalized to that of the nematode housekeeping gene *β-actin-1* and *EF-1α* with three biological replicates each (*n*=3). In (B), the transcript levels of each gene were normalized to that of the nematode housekeeping gene β-actin-1 with two biological replicates each (*n*=2). Data are presented as the mean ± SE. Each biological replicate contained a pool of hundreds of small root segments with infection sites for each post infection time point. dai, days after inoculation.

### *In situ* hybridization localizes *MigPSY* transcripts to the esophageal gland region of infective juveniles

We used *in situ* hybridization to localize *MigPSY2* transcripts in pre-parasitic J2s of *M. javanica* (**Figure 4A and 4B**) *and MigPSY3* transcripts in pre-parasitic J2s of *M. javanica* (**Figure 4C and 4D**) and *M. incognita* (**Figure 4E and 4F**). We detected a specific signal limited to the subventral esophageal glands when hybridized with a DIG-labeled antisense probe for both *MigPSY2* and *MigPSY3* (**Figure 4A, 4C and 4E**). We observed no hybridization when using a DIG-labeled sense probe as negative control for both species (**Figure 4B, 4D and 4F**). The subventral esophageal gland cells are thought to be the primary route by which secretions are delivered into the host during early stages of parasitism and have been shown to be most active at this stage (Hussey, 1989). Based on these observations, we hypothesize that MigPSYs are produced and sulfated in nematode subventral glands from where they are released into the host.

**Figure 4.**
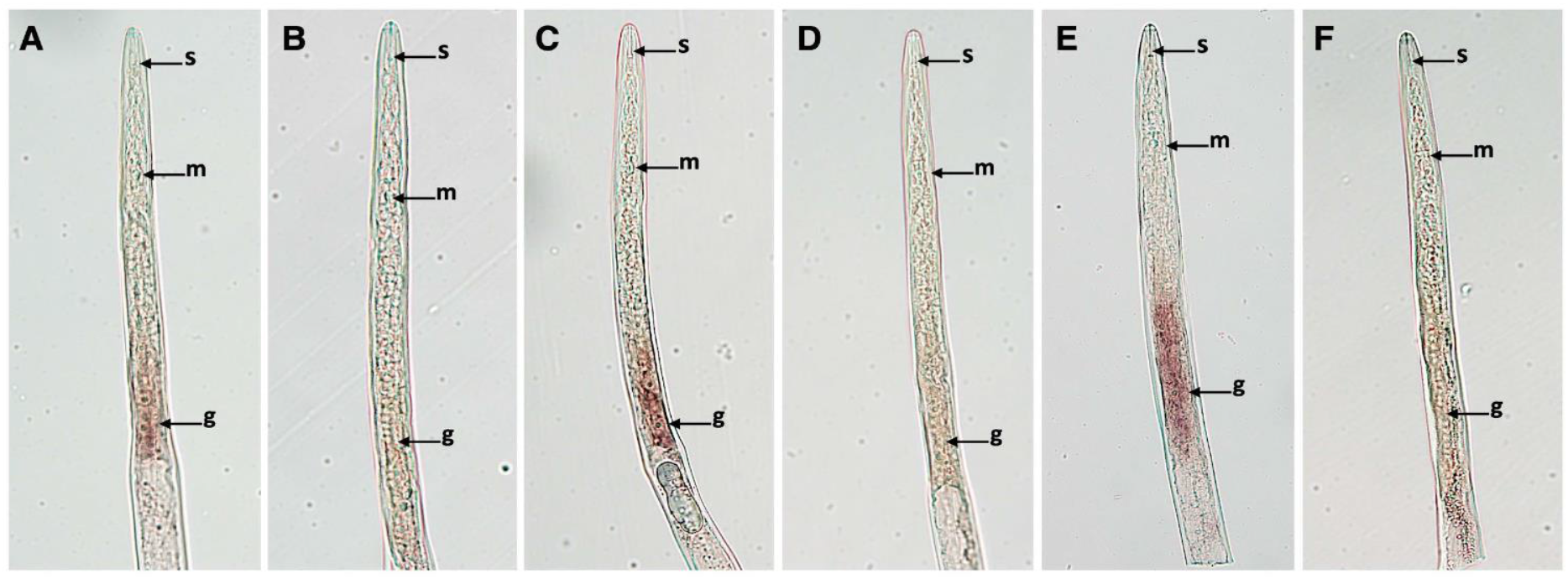
*In situ* hybridization of MigPSY transcripts in *Meloidogyne javanica and M. incognita* infective juveniles. (**A**) Positive detection of *MigPSY2* transcripts by an antisense DIG-labeled DNA probe in the esophageal gland region of *M. javanica*. (**C, E**) Positive detection of *MigPSY3* transcripts by an antisense DIG-labeled DNA probe in the esophageal gland region of *M. javanica* (C) and *M. incognita* (E). (**B, D, F**) As negative control, sense DIG-labeled DNA probes were used in *M. javanica* (**B, D**) and *M. incognita* (F) showing no positive labeling. Abbreviations: DIG, digoxigenin; s, stylet; m, metacorpus; g, glands. Procedure was repeated three times independently. Representative images are shown.

### Reduced expression of *MigPSYs* in infective J2 leads to fewer galls

To investigate the possible roles of MigPSY in parasitism, we soaked pre-parasitic J2 of *M. javanica* in a short interfering RNA (siRNA) targeting all *MigPSY*s present in *M. javanica*. A siRNA targeting the green fluorescent protein (*GFP*) gene was used as a control treatment. RT-qPCR showed that relative *MigPSY* transcript levels were lower in J2s treated with the *MigPSY* siRNA compared to control J2s (**Figure 5A**). We inoculated rice plants with siRNA-treated nematodes and examined their roots at 30 dai. We determined that the average number of galls on plants inoculated with J2s soaked in siRNA targeting *MigPSYs* was significantly lower than those inoculated with control J2s (**Figure 5B**). Following fixation of the roots and staining with acid fuchsin, we counted the total number of nematodes within the roots and found no differences between roots inoculated with J2s soaked in siRNA targeting *MigPSYs* and control roots (**Figure 5C**) suggesting that the reduced *MigPSY* transcript levels did not affect attraction to roots or infection. However, plants inoculated with infective J2s soaked in siRNA targeting *MigPSYs* had significantly fewer females with egg masses. Taken together, these results indicate a role for MigPSYs in nematode gall formation and in development of J2s into egg-laying females.

**Figure 5.**
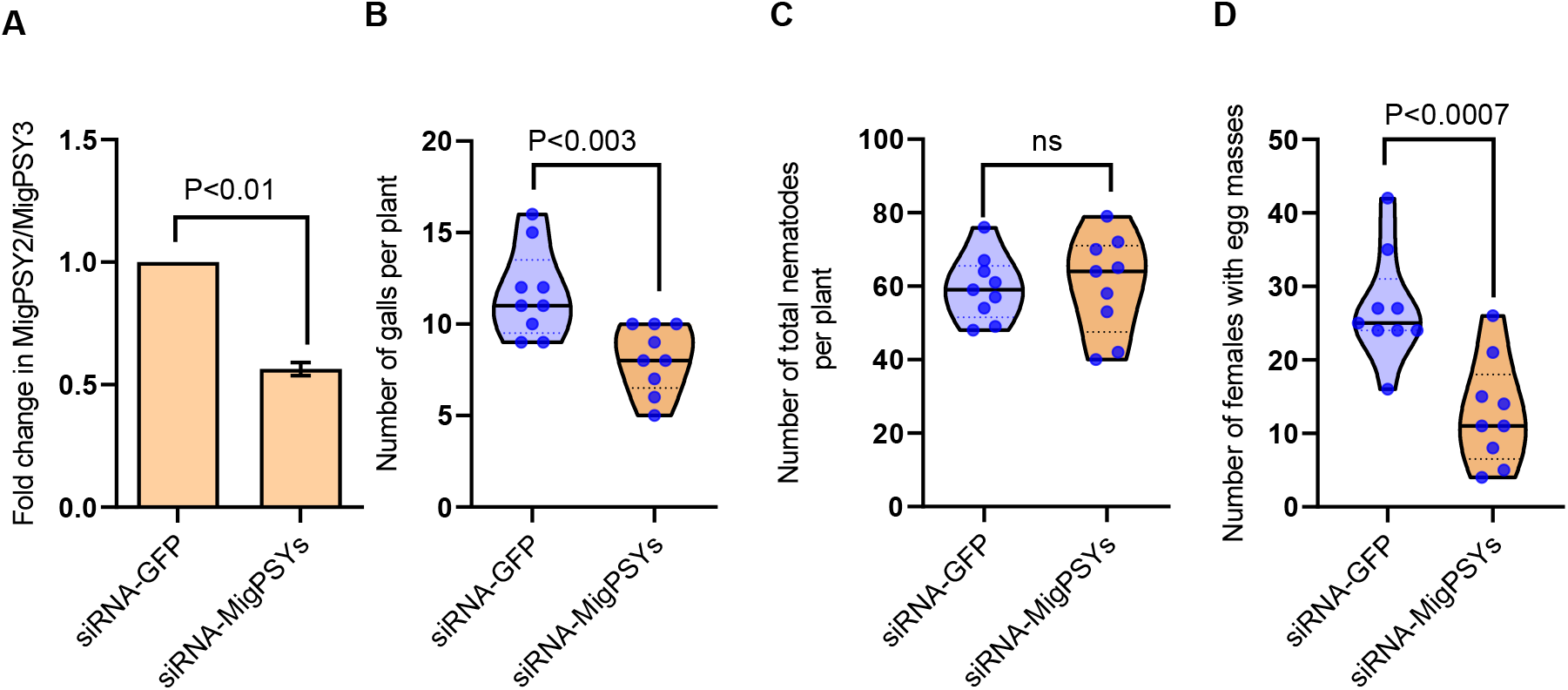
MigPSYs have a role in root-knot nematode parasitism. (**A**) Relative transcript abundance of *MigPSYs* in pre-parasitic J2s soaked in siRNA targeting either *MigPSYs* or *GFP*. Bars represent mean ± SEM from three independent experiments (*n*=3). Significant differences were assessed using Student’s *t*-tests (two-sided; *p*<0.05). (**B**) Number of galls present per rice root system infected with J2s soaked in siRNA targeting either *MigPSYs* or *GFP* at 30 dai. (**C**) Number of nematodes per rice root system infected with J2s soaked in siRNA targeting either *MigPSYs* or *GFP* at 30 dai. (**D**) Average number of females with egg masses per rice root system infected with J2s soaked in siRNA targeting either *MigPSYs* or *GFP* at 30 dai. In (**B-D**), approximately 1,000 *M. javanica* infective juveniles were inoculated onto 14-day-old rice seedlings. Experiments were performed three times independently with similar outcome. Data from one experiment are shown. Significant differences were assessed using Student’s *t*-tests (two-sided; *P*<0.05). Source data from all three experiment is included as supplementary file. For all violin plots, the center line is the median; dashed lines are upper and lower quartiles; the width of the violin represents the frequency; and points are the individual data points. MigPSY, *Meloidogyne incognita* group plant peptide containing sulfated tyrosine; siRNA, small-interfering RNA; J2, pre-parasitic second-stage juvenile; GFP, green fluorescent protein; dai, days after inoculation.

### Conclusions

In the present study, we report the identification and characterization of PSY peptides from root-knot nematodes. While RKN and *Xanthamonas* spp. have completely different life styles and parasitize different parts of the plant, both are biotrophic pathogens and induce hypertrophy of the host tissues. It is therefore likely that both RKN and *Xoo* employ PSY mimics to usurp related processes and modify the host environment to facilitate parasitism. However, the peptide sequences and expression patterns have likely been optimized to benefit each pathogen. Intriguingly, the conserved domain of MigPSY3 has a two-amino-acid insertion that to best of our knowledge is not present in any known plant PSY. The truncated synthetic peptide MigPSY3^-C^ did not stimulate root elongation in our Arabidopsis assay as strongly as the similarly truncated peptides for MigPSY1 or MigPSY2 suggesting that the 2 aa insertion may alter binding to the plant receptor. However, MigPSY3 is the most widespread and conserved among the closely related MIG species, suggesting that it has a valuable role in this highly successful group of parasites. Understanding the contribution of MigPSYs to gall formation and nematode reproduction will require additional work to identify post translational modifications, timing and targeting of delivery of these peptides in the host.

## Methods

### Gene identification and characterization

We used the following databases to identify nematode genes and transcripts described in this work: https://www6.inrae.fr/meloidogyne_incognita; https://parasite.wormbase.org/index.html. Translation products were characterized for signal peptide sequences using SignalP-6.0 (https://services.healthtech.dtu.dk/service.php?SignalP), and the presence of any transmembrane domain was assessed by DeepTMHMM analysis https://doi.org/10.1101/2022.04.08.487609).

Sequence logos were generated using WebLogo (http://weblogo.berkeley.edu/).

### Plant growth and nematode inoculation

Pure cultures of *Meloidogyne javanica* strain VW4 were maintained on *Solanum lycopersicum* cv. Momar Verte plants grown in sand in a greenhouse (Wang et al., 2009).

Nematode eggs were collected from three-month-old cultures, and pre-parasitic J2s were hatched as described (Branch et al., 2004) with minor modifications. Seeds of rice (*Oryza sativa* L.) subsp. *japonica* ‘Kitaake’ and tomato (*Solanum lycopersicum*) cultivar ‘Moneymaker’ were surface sterilized in 70% (v/v) ethanol and 4% sodium hypochlorite for 30 min and germinated on wet filter paper at 28°C in darkness for 3 days. Seedlings were then transplanted and grown in Sand Absorbent Polymer (SAP) substrate in a growth chamber (26°C; 12 h light/12 h dark regime). Plants were fertilized with Hoagland solution three times a week (Reversat et al., 1999). Two-week-old rice and tomato seedlings were inoculated with 2,000 J2s of *M. javanica* or mock-inoculated with water as a control. Nematode infection was evaluated 30 dai as described (Yimer et al., 2018).

### RNA extraction, cDNA synthesis, and qRT-PCR

For gene expression analysis of *MigPSYs* during *M. javanica* parasitism of tomato and rice, plants were grown in sand and inoculated with nematodes as above. The entire root (for rice) and root tips or nematode galls (for tomato) was harvested at 2, 4, 8, and 21 dai with *M. javanica*. Samples were immediately frozen in liquid nitrogen and stored at –80°C. Total RNA was extracted using an RNeasy Plant Mini Kit (Qiagen, USA) following the manufacturer’s instructions. RNA concentration and purity were measured using a NanoDrop OneC Microvolume UV-Vis Spectrophotometer (Thermo Scientific, USA). TURBO DNase treatment was carried out to remove genomic DNA from total RNA samples (TURBO DNA-free Kit™, Ambion, USA). One-step RT-qPCR was performed using an iTaq Universal SYBR Green One-Step RT-qPCR Kit (BIO-RAD). The reaction was performed in a total volume of 20 μL by mixing 10 μL iTaq universal SYBR Green reaction, 0.25 μL of iScript reverse transcriptase, 1 μL of each primer (10 μM), 5.75 μL nuclease-free water, and 1.5 μL (80 ng μL^−1^) DNase treated RNA. The reaction was carried out with an Applied Biosystem QuantStudio 3 Real-Time PCR System using the following conditions: reverse transcription at 50°C for 10 min, polymerase activation at 95°C for 1 min, followed by 40 cycles of denaturation at 95°C for 15 s and annealing/extension and plate reading at 60°C for 60 s. A melting curve analysis was conducted by gradually increasing the temperature to 95°C. The expression analysis was performed in triplicate using three independent biological samples, consisting of a pool of eight plants each. The transcript levels of *MigPSY*s were normalized to those of the nematode housekeeping gene *β-actin-1* and *EF-1α* in the tomato experiment and *β-actin-1* in the rice experiment (Iberkleid et al., 2013, Painter & Lambert, 2003). Relative gene expression data were computed according to Pfaffl (Pfaffl, 2001).

### Root growth assays

*Arabidopsis thaliana tpst-1* mutant seeds were surface-sterilized for 10 min with 70% (v/v) ethanol and then stratified in 0.1% agarose in the dark (4°C) for 3-4 days. Solid nutrient media plates were prepared with 1X Murashige and Skoog (MS) medium with vitamins (MSP09; Caisson Labs (East Smithfield, UT, USA)), 1% sucrose, pH 5.7, 0.3% Gelzan (G024; Caisson Labs). Tyrosine sulfated peptides (listed in Supplementary Table S3) were synthesized by the Protein Chemistry Facility at the Gregor Mendel Institute and resuspended in double-distilled water. Synthetic peptide (or water for mock treatments) was added to the medium to a final concentration of 100 nM just before pouring into a 100 mm × 15 mm Petri dish. Seeds were distributed equally on the surface of the medium along a row 2-3 cm from the upper edge of the Petri dish and the lids were secured with Micropore surgical tape. Plates were incubated vertically in a 21°C chamber with 16-h-light/8-h-dark photoperiod. Seedlings with delayed germination were marked after 3 days and were not included in the analysis. Root lengths (mm) were measured 9 days after sowing using ImageJ software (version 1.53, National Institutes of Health).

### *In situ* hybridization

Total RNA was extracted from pre-parasitic J2s of both *M. javanica* and *M. incognita* using RNeasy Plant Mini Kit (Qiagen) and the first-strand cDNA synthesized with High-Capacity cDNA Reverse Transcription Kits (Applied Biosystems, Thermo Scientific). The cDNA template was then used to PCR amplify 79 bp in the CDs region of *MigPSY2* (*M. javanica*) and 81 and 88 bp in the CDS region of *MigPSY3 (M. javanica* and *M. incognita)*. The primers used for probe synthesis are listed in Supplementary Table S4. The *in-situ* hybridization protocol was adopted from (Gao et al., 2003). A sense (negative control) and anti-sense single-strand cDNA probe were synthesized in two independent asymmetric PCR reactions with digoxigenin (DIG) labeling mix (1mM dATP, 1mM dCTP, 1mM dGTP, 0.65mM dTTP, 0.35mM DIG-dUTP; Roche, CA, USA). Pre-parasitic J2s of *M. javanica* and *M. incognita* were fixed in 2% paraformaldehyde in M9 buffer for 18 h at 4°C, followed by 4 h of incubation at room temperature. The fixed pre-parasitic J2s were mechanically cut into fragments and enzymatically permeabilized with proteinase K (0.5 mg/mL) solution (Roche, Germany). Hybridization with sense (negative control) or anti-sense dig-labeled probes was performed in separate nematode samples overnight at 50°C. Hybridized probes were detected by anti-DIG antibodies conjugated with alkaline phosphatase enzyme (anti-DIG-AP) (Roche, Germany). Photographs were captured using an Olympus BX51 compound microscope using a DP74 Olympus camera.

### Gene silencing

*MigPSY* transcript levels were knocked down using an siRNA as described previously (Siddique S et al., 2021). Briefly, a 21-nt long target sequence in the *MigPSY3* mRNA beginning with an AA dinucleotide was identified. Sense and antisense oligonucleotides for the 21-nt target sequence were obtained in which the U’s were substituted with T’s. The 8-nt long sequence (5’-CCTGTCTC-3’) complementary to the T7 promoter primer was added to the 3’ ends of both the sense and antisense oligonucleotides. Afterwards, a double-stranded RNA (dsRNA) was synthesized from the sense and antisense oligonucleotides using a Silencer® siRNA Construction Kit (Cat. Nr. AM1620) according to the manufacturer’s instructions. The siRNA targeting *eGFP* was used as a control. siRNA yields were measured using a NanoDrop OneC Microvolume UV-Vis Spectrophotometer (Thermo Scientific, USA). Approximately 10,000 freshly hatched pre-parasitic J2s were incubated in 50 μL solution containing 200 ng/uL of siRNA, 3 mM spermidine (Sigma-Aldrich, USA), and 50 mM octopamine (Sigma-Aldrich, USA). After a 24-h incubation at 26°C in the dark, nematodes were washed with sterile water several times. J2s were then incubated for another 24 h in sterile water before sampling for RNA extraction. Total RNA was extracted from J2s subjected to siRNA targeting either *MigPSY* or *GFP*, and one-step RT-qPCR was performed as described above to analyze the *MigPSY* transcript levels. For infection assay, rice seeds were germinated and grown in sand as described above. The viability of siRNA-treated nematodes was observed under a microscope before inoculation. Two-week-old rice seedlings were inoculated with 1000 J2s. Two days post-inoculation, plants were transferred to a hydroponic system to synchronize the infection. In brief, plants were carefully uprooted from the sand substrate, washed under tap water to remove the adhering sand particles, and transferred to 1/4 strength Hoagland’s solution. Plants were maintained under hydroponic conditions for 28 days, and nematode infection was evaluated 30 dai with *M. javanica*. Roots were stained with acid fuchsin to stain nematodes. Nematodes inside roots were enumerated by microscopic examination. Statistical analysis and graphical representations were performed using GraphPad Prism software version 8.3.0.

## Acknowledgements

The work was supported by NSF IOS 1954929. We thank Youssef Belkhadir (Gregor Mendel Institute of Molecular Plant Biology, Vienna, Austria) for providing sulfated peptides used in this work.

