## Supplementary Data for "Root-knot nematodes produce functional mimics of tyrosine-sulfated plant peptides"

**Supplementary materials**

**Table S1. PSY-like peptides encoded in MIG species.** Peptides followed by the same letter are identical in sequence.

| **cDNA designation** | **MigPSY type** | **# Amino acids in encoded peptide** | **Predicted peptide length after signal cleavage^1^** |
| --- | --- | --- | --- |
| M.Arenaria_Scaff18626g092470 | PSY1 | 51 | 21 |
| M.Arenaria_Scaff5652g054995 | PSY2 | 51 | 21 |
| M.Javanica_Scaff4941g037068 | PSY2 | 51 | 21 |
| M.Arenaria_Scaff13461g082170 | PSY3^a^ | 50 | 23 |
| M.Arenaria_Scaff5456g053959 | PSY3^b^ | 50 | 23 |
| M.Arenaria_Scaff20106g094880 | PSY3^c^ | 50 | 23 |
| Minc3s01748g26028 | PSY3^a^ | 50 | 23 |
| Minc3s07420g41102 | PSY3^b^ | 50 | 23 |
| Minc3s02310g29383 | PSY3^c^ | 51 | 24 |
| M.Javanica_Scaff5264g038601 | PSY3^a^ | 50 | 23 |
| M.Javanica_Scaff14581g070241 | PSY3^b^ | 50 | 23 |

**Table S2. Amino acid sequences for PSY-like peptides encoded in MIG species.** Underlined letters represent highly a conserved N-terminal signal peptide. Letters in red represent a conserved tyrosine residue and letters in blue represent a PSY-like domain with additional C-terminal GGGR sequence.

| MaPSY1: | MNTSLLFNFVTLSIIYVILYLSFAEAYTINDYG--GPSANDRHDPLKGL-GGGR |
| --- | --- |
| MaPSY2: | MNTSLLFNFVTLSIIYVILYLSFAEAYTINDYP--ETGPNHHHDPPKRL-GGGR |
| MjPSY2: | MNTSLLFNFVTLSIIYVILYLSFAEAYTINDYP--ETGPNHHHDPPKRL-GGGR |
| MaPSY3^a^: | MNTSILFNFVTLSIIYVILYLSFTEA---SDYGSRSPGANDAHDPKKQL-GGGR |
| MaPSY3^b^: | MNTSILFNFVTLSIIYVILYLSFAEA---LDYGSRSPGANDAHDPKKQL-GGGR |
| MaPSY3^c^: | MNTSILFNFVTLSIIYVILYLSFAEA---LDYGSRSPGANDAHDPSKQL-GGGR |
| MiPSY3^a^: | MNTSLLFNFVTLSIIYVILYLSFAEA---SDYGSRSPGANDAHDPKKQLRGGGR |
| MiPSY3^b^: | MNTSILFNFVTLSIIYVILYLSFTEA---SDYGSRSPGANDAHDPKKQL-GGGR |
| MiPSY3^c^: | MNTSILFNFVTLSIIYVILYLSFTEA---SDYGSRSPGANDAHDPKKQL-GGGR |
| MjPSY3^a^: | MNTSILFNFVTLSIIYVILYLSFAEA---LDYGSRSPGANDAHDPSKQL-GGGR |
| MjPSY3^b^: | MNTSILFNFVTLSIIYVILYLSFTEA---SDYGSRSPGANDAHDPKKQL-GGGR |

**Table S3: Synthetic peptide sequences.**

| **Peptide** | **Sequence** |
| --- | --- |
| AtPSY1 | DY(SO_3_)GDPSANPKHDPGVPPS |
| RaxX21 | HVGGGDY(SO_3_)PPPGANPKHDPPPR |
| RaxX13^-C^ | DY(SO_3_)PPPGANPKHDP |
| MigPSY1 | DY(SO_3_)GGPSANDRHDPLKGLGGGR |
| MigPSY1^-C^ | DY(SO_3_)GGPSANDRHDP |
| MigPSY2 | DY(SO_3_)PETGPNHHHDPPKRLGGGR |
| MigPSY2^-C^ | DY(SO_3_)PETGPNHHHDP |
| MigPSY3 | DY(SO_3_)GSRSPGANDAHDPKKQLGGGR |
| MigPSY3^-C^ | DY(SO_3_)GSRSPGANDAHDP |

**Table S4: Primer sequences used in this study.**

| **Primer name** | **Sequence** |
| --- | --- |
| MjPSY3_ISH_F | CGCTGAAGCATTAGATTATGGA |
| MjPSY3_ISH_R | CGTCCTCCTCCTAATTGTTTTG |
| MiPSY3_ISH_F | CATTCGCTGAAGCATCAGATT |
| MiPSY3_ISH_F | CGTCCTCCTCCTCTTAATTGTT |
| MjPSY2_ISH_F2 | CATTCGCTGAAGCATACACC |
| MjPSY2_ISH_R | CCTAATCGTTTTGGTGGATCA |

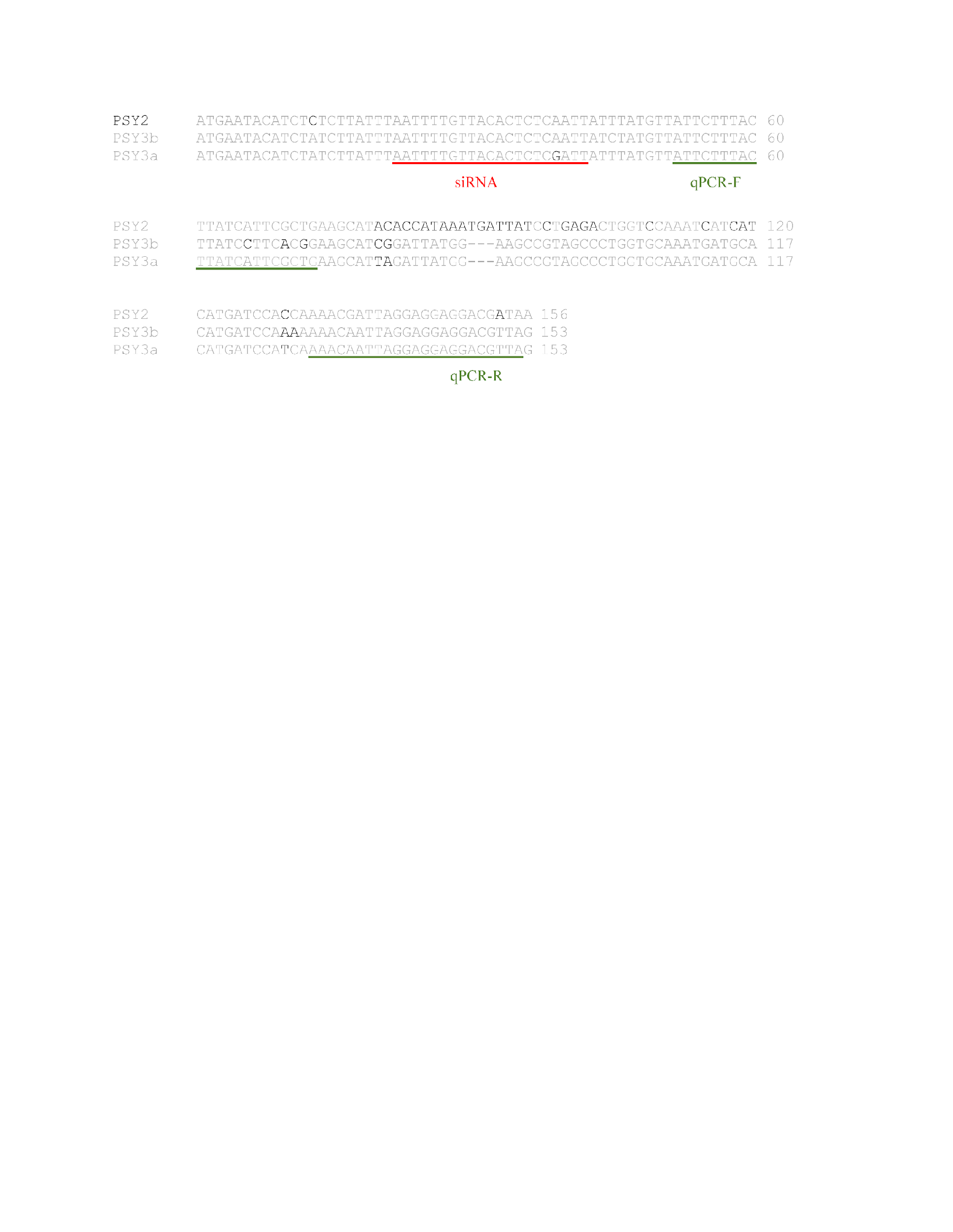

**Supplementary Figure 1.** CDs alignment of MigPSYs from *Meloidogyne javanica.* Highlighted regions are siRNA target site (red) and qPCR primer sites (green).
